# Tubewell use protects against rotavirus infection during the monsoons in an urban setting

**DOI:** 10.1101/630855

**Authors:** Pamela P. Martinez, Ayesha Mahmud, Mohammad Yunus, A.S.G Faruque, Tahmeed Ahmed, Mercedes Pascual, Caroline O. Buckee

## Abstract

Rotavirus, a diarrheal pathogen spread via fecal-oral transmission, is typically characterized by a winter incidence peak in most countries. Unlike for cholera and other water-borne infections, the role of environmental and socioeconomic factors on the spatial variation of rotavirus seasonality remains unclear. Here, we analyze their association with rotavirus seasonality, specifically the odds of monsoon cases, across 46 locations from 2001 to 2012 in Dhaka. Drinking water from tubewells, compared to other sources, has a clear protective effect against cases during the monsoon, when flooding and water contamination are more likely. This finding supports a significant environmental component of transmission.

## Introduction/Background

Rotavirus is the most common cause of diarrheal deaths among children below the age of 5 worldwide, causing more than 215,000 deaths in 2013 [1]. Globally, rotavirus exhibits significant heterogeneity in seasonality. Highest incidence typically occurs during cooler and dryer seasons in most countries [2, 3]. In some regions, where the disease tends to have a more endemic transmission, an additional peak occurs during the summer or monsoon season, for reasons that are not well defined [4, 5]. Bangladesh is an example of a country with high incidence all year round, particularly in Dhaka the capital city, where high temperatures, low humidity, and flooding events have been associated with an increased risk for rotavirus [6]. Current dogma suggests that person-to-person contact is the dominant mode of transmission of the virus, although it has been shown to persist in water [7] and to respond to climatic changes [8]. Dhaka has more than 18 million inhabitants and a population growth of about 4% annually, making it one of the densest and fastest growing megacities in the world. Heterogeneities in urbanization within Dhaka have been previously shown to affect the sensitivity of diarrheal diseases to climate forcing [8, 9].

Here, we focus on seasonality and examine its association with several socio-economic and environmental variables. We leverage spatial variation in seasonal patterns across 11 years of data, to identify the external factors that are associated with disease transmission at different times of the year. This is of particular relevance in places like Bangladesh, where vaccine effectiveness is still low [10] and rotavirus incidence remains high.

## Methods

### Data

A total of 8481 confirmed rotavirus cases between 2001 and 2016 were obtained from the Dhaka Hospital, through the ongoing surveillance program of the International Centre for Diarrheal Disease Research, Bangladesh (icddr,b). Stool samples were collected to determine the presence of enteric pathogens using an enzyme-linked immunosorbent assay, for every 50^th^ patient attending the hospital for treatment of diarrhea. Each case in the dataset includes the residence location of the patient. Since administrative boundaries within the Dhaka Metropolitan Area changed over the time period of our analysis, we aggregated the data for newer, smaller administrative units to match the 2001 boundary definitions (Figure S1). This ensures that the spatial patterns are consistent across the period of analysis. For a subset of patients (around 11%) who tested positive for rotavirus between January 2001 and May 2012 (658 of the 5833 confirmed cases for this period), we have additional demographic and socio-economic information from a survey administered to the patient.

### Statistical analysis

We computed the Moran’s index (Moran’s I) to examine spatial patterns in rotavirus seasonality. The global Moran’s I was used to measure the degree of spatial autocorrelation in the fraction of cases occurring during the monsoon across all locations in our data; the local Moran’s I was computed to identify local spatial clusters. The Moran statistics were calculated assuming neighboring locations had equal weight. We used a contiguity neighbor definition based on neighbors sharing at least one polygon vertex, but our results are robust to using a distance-based neighbor definition (Figure S2). We computed pseudo p-values for the Moran statistics by comparing the observed values to a sampling distribution of Moran’s I values generated using a random distribution assumption.

We used a generalized linear mixed effects model to identify the relative importance of demographic, economic, and environmental factors on the likelihood of cases occurring during the monsoon season relative to the rest of the year. The outcome of interest is a binary variable indicating whether or not a case occurred during the monsoon season. Specifically, we modeled the log odds of the outcome variable as a linear combination of a set of independent variables and a random effect for each location. The full model specification is:

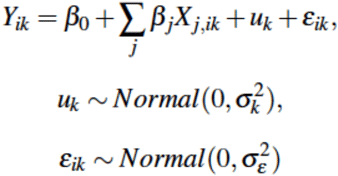

where *Y_ik_* is the log-odds of a case occurring in the monsoon season for individual *i* in location *k*; *β_0_* is the intercept (grand mean across all individuals and locations); *X_j,ik_* are a set of individual-level independent variables; *β_j_* are the corresponding estimated fixed effects; *u_k_* are the estimated random effects (location-specific deviation from the grand mean); and *ε_ik_* are the model residuals. The random effects are assumed to be normally distributed with mean zero and variance, *σ^2^_k_*.

We tested three sets of fixed effects at the individual level: (1) demographic factors (age and sex); (2) environmental factors (use of tubewell, defecating in the open, and treatment of drinking water); and (3) socioeconomic factors (father’s income and the type of material the roof of the individuals’ houses are constructed with). All independent variables were normalized to have mean zero and standard deviation one. We compared models containing combinations of the different sets of independent variables (Table S1), and report results for the full model including all variables and for the model with the lowest Akaike Information Criteria (AIC). The 95% confidence intervals (CI) for the fixed effect estimates were computed using simulations from the posterior distribution.

## Results

We first characterized the spatial heterogeneity of rotavirus seasonality by classifying cases into ‘monsoon cases’ (Apr–Sep) and ‘winter cases’ (Jan–Mar, Oct–Dec), aggregated for each location. The average annual fraction of rotavirus cases during the monsoon season exhibits a strong latitudinal gradient across the locations in our data, with locations in the south and in the Dhaka Metropolitan Area experiencing a higher percentage of cases during this season (Figure 1A). We detected significant spatial autocorrelation in rotavirus seasonality (global Moran’s I = 0.37, p-value < 0.05), and the existence of spatial clustering, indicated by a significant local Moran’s I statistic, particularly in the north of the city (Figure 1B). The clustering of locations in the north captures locations that have a low fraction of cases during the monsoons surrounded by locations whose fraction is also low.

**Figure 1.**
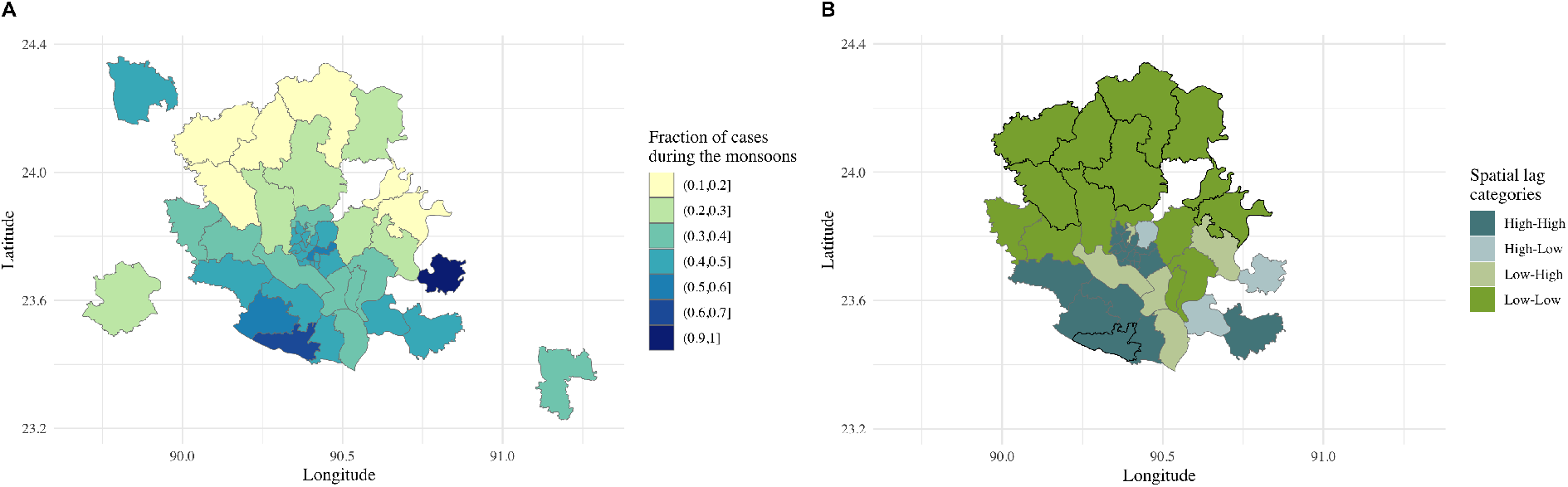
Rotavirus seasonality. (A) Average annual fraction of rotavirus cases occurring during the monsoon season for each location, based on 16 years of data. (B) Spatial clusters in rotavirus seasonality identified using local Moran’s I. Locations with a significant (pseudo p-value < 0.05) local Moran’s I statistic are shown with black borders; colors indicate the type of spatial classification. High-High (Low-Low) indicates locations with a high (low) fraction of cases during the monsoons that are surrounded by locations with a high (low) fraction; High-Low (Low-High) indicates locations with a high (low) fraction of cases during the monsoons that are surrounded by a low (high) fraction.

To characterize the drivers of this spatial heterogeneity in the seasonality of the aggregated data, we analyzed the individual level survey data. The spatial patterns in winter (Figure 2A) and monsoon (Figure 2B) incidence in these survey data are consistent with the pattern for all reported cases (Figure S3), indicating that the subset data is representative of the overall incidence data. When we model rotavirus seasonality as a function of demographic, environmental, and socioeconomic factors, at the individual-level, we find a statistically significant association between reported tubewell use and the odds of a case occurring during the monsoon season versus the rest of the year (Figure 2C). Specifically, we find that in the selected model (lowest AIC) individuals who obtained their drinking water primarily from a tubewell, rather than tap, pond, or other sources, exhibit decreased odds of infection during the monsoons. The odds were 0.47 times lower (95% CI, 0.31–0.71), holding all else (including the random effects) fixed. This change corresponds to a 53%decrease in the odds of experiencing a monsoon case for those reporting tubewell use compared to those relying on other sources for their drinking water. We find no other statistically significant associations.

**Figure 2.**
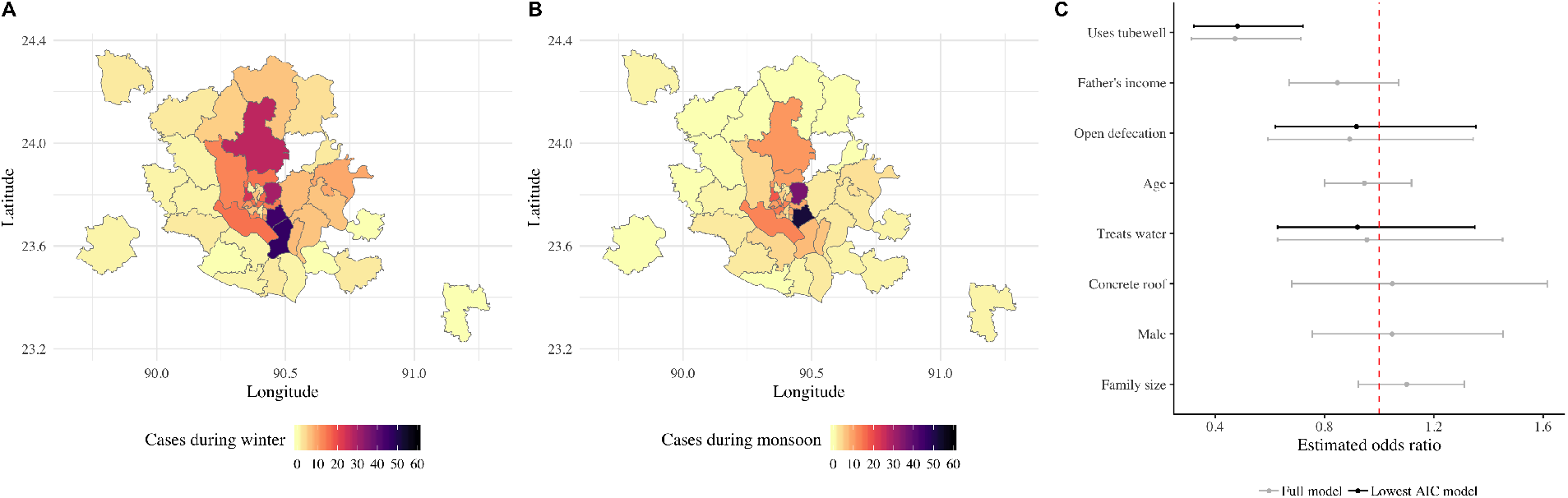
Spatial distribution of cases in the individual-level survey data that occurred during the winter (A) and the monsoon (B) seasons. (C) Estimated fixed effects (and 95% CI) for the full model (in grey) and for the lowest AIC model (in black). The estimated fixed effects represent the estimated association between each of the independent variables and the odds of a case occurring during the monsoons, holding all else fixed (including the random effects). The red dashed line indicates an odds ratio of 1.

## Discussion

Our findings show that drinking water from tubewells, which are less likely to be contaminated during flooding events, is protective against rotavirus during the monsoon season. While rotavirus transmission is typically considered to occur via direct person-to-person contact, this result highlights the importance of environmental transmission via water in Dhaka. This is consistent with recent studies showing that rotavirus transmission is sensitive to interannual variation in flooding [8], and that waterborne transmission alone is sufficient to spread the virus between hydrologically connected locations [7]. The use of tubewells has also been shown to provide a protective effect against other diarrheal diseases in Bangladesh, such as cholera [11, 12] and shigellosis [12], especially in areas that are not protected from flooding [13]. Tubewells are less likely to be contaminated during flooding events as they source water from deeper underground compared to other water supply systems. The consequences of a possible environmental reservoir are unclear, especially in Bangladesh, where recent shifts in monsoon rainfall have been observed across different regions [14, 15]. An increase in heavy precipitation has the potential to trigger flooding events, and in doing so, increase the incidence of rotavirus and other water-borne diarrheal diseases.

There are several limitations that should be noted. Our analyses compared individuals with confirmed rotavirus infection at different times of the year. Since there were no control group without the disease, we were unable to test whether tubewell use is protective against rotavirus in general. We also do not provide a mechanistic explanation for the spatial gradient observed in rotavirus seasonality at the aggregate-level, or for the differences observed between winter and monsoon incidence. Thus, we cannot rule out potential unobserved confounders that could also explain the seasonal pattern described here. More detailed spatiotemporal data, with a well-defined control group, is needed in order to address some of these limitations.

In conclusion, our findings support the hypothesis that consumption of clean water plays an important role in protecting against rotavirus infection, particularly during the monsoon season, when flooding can contaminate surface water. The potential transmission of rotavirus through water should be recognized when planning control interventions. This is especially important in highly populated regions of developing countries where endemic transmission may be modulated seasonally by the existence of environmental reservoirs.

## Acknowledgments

The case data used in this paper were collected with the support of the ICDDR,B and its donors, who provide unrestricted support to the ICDDR,B for its operation and research. Current donors providing unrestricted support include the Government of the People’s Republic of Bangladesh; the Department of Foreign Affairs, Trade, and Development Canada; the Swedish International Development Cooperation Agency; and the Department for International Development (UK Aid). We thank these donors for their support and commitment to the ICDDR,B’s research efforts. P.P.M. was funded by NIH grant R01 AI048935. A.M. acknowledges funding from The Rockefeller Foundation Planetary Health Fellowship. C.O.B was supported by NIH grant R35 GM124715-02.

## Supplementary figures

**Figure S1.**
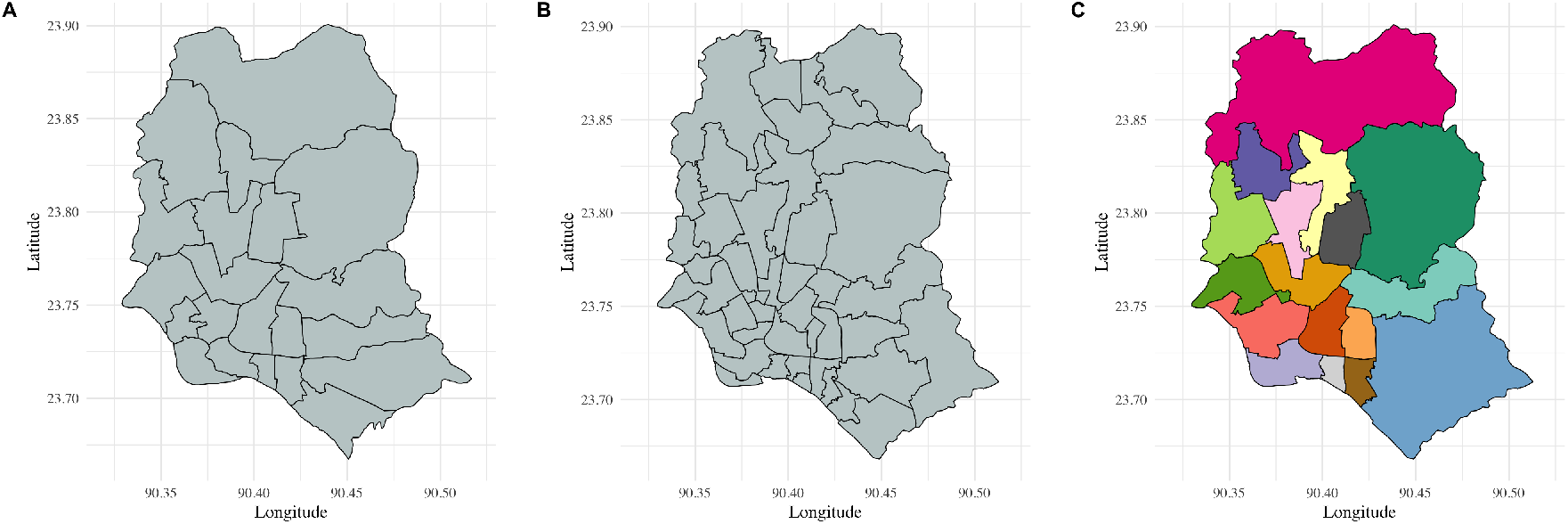
Administrative subdivisions and location aggregation for the Dhaka Metropolitan Area. (A) Administrative subdivision boundaries established in 2001. (B) Administrative subdivision boundaries established in 2011. (C) Aggregated administrative subdivisions used for consistency across years. Each color represents a different location used in the current study.

**Figure S2.**
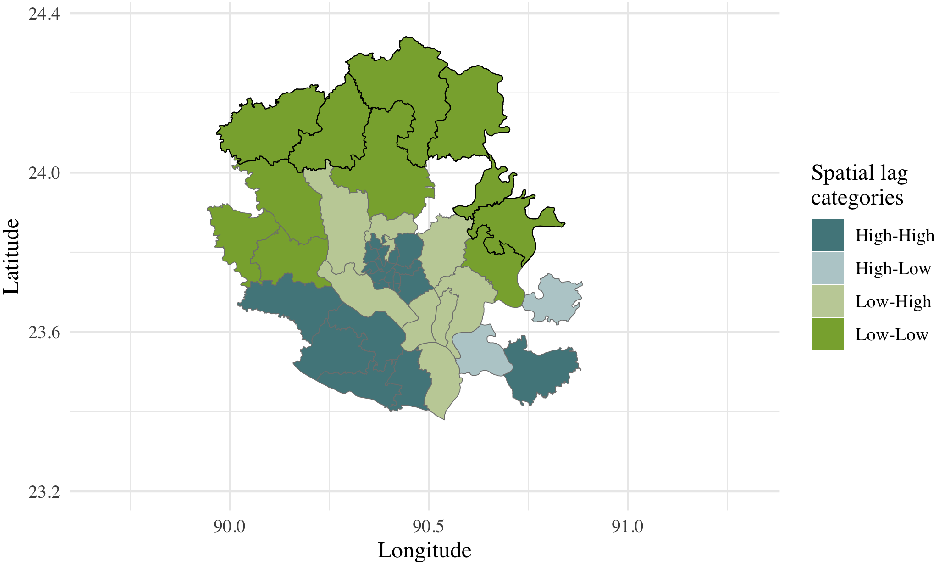
Spatial clusters in rotavirus seasonality identified using local Moran’s I and a distance-based neighbor definition. Locations with a significant local Moran’s I statistic are shown with black borders.

**Figure S3.**
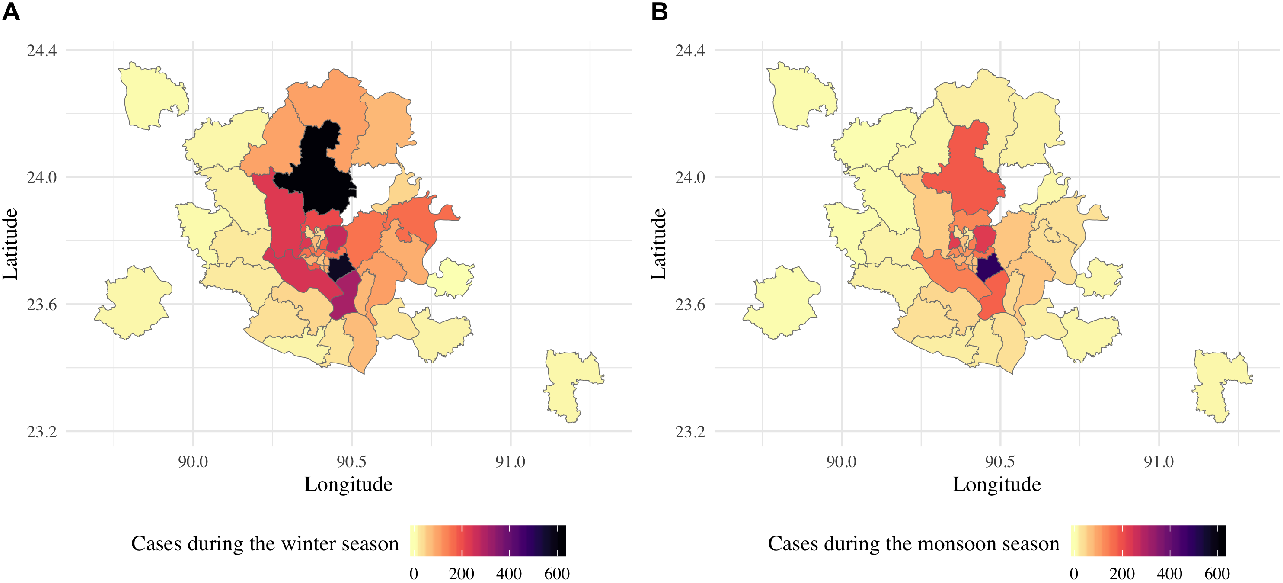
Spatial distribution of cases in the aggregated data set that occurred during the winter (A) and the monsoon (B) seasons.

**Table S1.** Estimated fixed effects for all the possible combinations of the different sets of independent variables (demographic, environmental, and socioeconomic factors).

|  | *Dependent variable: Log odds of a case occurring in the monsoon season* |  |  |  |  |  |  |
| --- | --- | --- | --- | --- | --- | --- | --- |
|  | (1) | (2) | (3) | (4) | (5) | (6) | (7) |
| Age | −0.063<br>(0.084) |  |  | −0.047<br>(0.085) | −0.072<br>(0.085) |  | −0.056<br>(0.085) |
| Male | 0.077<br>(0.170) |  |  | 0.073<br>(0.171) | 0.065<br>(0.170) |  | 0.047<br>(0.171) |
| Uses Tubewell |  | −0.745***<br>(0.213) |  |  | −0.750***<br>(0.214) | −0.747***<br>(0.215) | −0.751***<br>(0.216) |
| Open defecation |  | −0.083<br>(0.203) |  |  | −0.091<br>(0.204) | −0.108<br>(0.209) | −0.112<br>(0.209) |
| Treats water |  | −0.065<br>(0.202) |  |  | −0.070<br>(0.202) | −0.040<br>(0.215) | −0.047<br>(0.215) |
| Family Size |  |  | 0.061<br>(0.089) | 0.058<br>(0.090) |  | 0.098<br>(0.089) | 0.096<br>(0.090) |
| Father's income |  |  | −0.172<br>(0.114) | −0.166<br>(0.116) |  | −0.171<br>(0.117) | −0.164<br>(0.120) |
| Concrete roof |  |  | 0.229<br>(0.202) | 0.231<br>(0.203) |  | 0.045<br>(0.222) | 0.046<br>(0.222) |
| Constant | −0.504***<br>(0.164) | −0.066<br>(0.188) | −0.523***<br>(0.137) | −0.569***<br>(0.176) | −0.100<br>(0.217) | −0.087<br>(0.196) | −0.110<br>(0.226) |
| Observations | 658 | 658 | 658 | 658 | 658 | 658 | 658 |
| Akaike Inf. Crit. | 880.547 | 868.678 | 879.836 | 883.354 | 871.793 | 871.436 | 874.924 |

*Note:* \*p<0.1; \*\*p<0.05; \*\*\*p<0.01

